# Inferring multi-locus selection in admixed populations

**DOI:** 10.1101/2023.05.15.540874

**Authors:** Nicolas M. Ayala, Russell Corbett-Detig

## Abstract

Admixture, the exchange of genetic information between distinct source populations, is thought to be a major source of adaptive novelty. Unlike mutation events, which periodically generate single alleles, admixture can introduce many selected alleles simultaneously. As such, the effects of linkage between selected alleles may be especially pronounced in admixed populations. However, existing tools for identifying selected mutations within admixed populations only account for selection at a single site, overlooking phenomena such as interference among proximal selected alleles. Here, we develop and extensively validate a method for identifying and quantifying the individual effects of multiple linked selected sites on a chromosome in admixed populations. Our approach numerically calculates the expected local ancestry landscape in an admixed population for a given multi-locus selection model, and then maximizes the likelihood of the model. After applying this method to admixed populations of *Drosophila melanogaster,* we found that the impacts between linked sites may be an important contributor to natural selection in admixed populations. Furthermore, for the situations we considered, the selection coefficients and number of selected sites are overestimated in analyses that do not consider the effects of linkage among selected sites. Our results imply that linkage among selected sites may be an important evolutionary force in admixed populations. This tool provides a powerful generalized method to investigate these crucial phenomena in diverse populations.

## Introduction

Admixture is one of the primary sources of selected alleles in natural populations [1–3]. For example, in *Helianthus* sunflowers, introgressed alleles enhanced herbivore resistance at a number of loci [4]. In the fish *Fundulus grandis,* recently introgressed alleles allow resistance to extreme pollution and environmental change [5]. In humans, introgression from archaic hominids is thought to have facilitated adaptation to a range of novel environments [6,7]. Similarly, admixture can contribute alleles that are not adapted to local environments (*e.g.,* as in [8]) or it may contribute haplotypes that are deleterious due to accumulation of weakly deleterious mutations during long term isolation in small populations [9], and are therefore purged by natural selection. Finally, in some cases admixed populations may contain mutations contributed by separate parental populations that have negative interactions thereby resulting in strong selection within the admixed populations [8,10]. Although the importance of selection on admixed ancestry, or adaptive introgression, is increasingly appreciated, generalized methods to accurately detect and quantify the impacts of natural selection from genome sequence data in admixed populations are in their infancy.

Admixture may be disproportionately likely to create circumstances where selected sites affect the evolutionary dynamics of other selected sites through linkages. The effects of multi-locus selection have been studied extensively in the context of populations along a geographic cline. Theory demonstrates that closely linked locally selected alleles can strongly reshape their expected frequencies across geographic clines (*e.g.*, [11–13]), and fixation probabilities in a continent-island model (*e.g.*, [14–17]). More generally, linkage among selected alleles can generate complex dynamics in clines that include combinations of loci involved in complex selection (*e.g.*, [18]). Although multi-locus selection in admixed populations generated through a single admixture event, sometimes called an “admixture pulse”, has been less extensively studied (although see *e.g.,* [9]), linkage among selected alleles should also impact evolutionary dynamics in such admixed populations. For example, if each ancestral population contributes distinct adaptive variants that are closely linked, their fixation could be impeded by Hill-Robertson interference [19,20]. Conversely, and of particular relevance to our application below, if a single population contributes linked adaptive variants, their collective allele frequency change could exceed expectations for single loci due to synergistic hitchhiking effects. For example, we might expect the latter in circumstances where one ancestral population has recently undergone polygenic adaptation for a trait that remains beneficial within the admixed population. It is therefore important to develop inference methods for detecting and quantifying the impacts of multiple selected alleles within admixed populations.

General frameworks for detecting selection within admixed populations have developed substantially in recent years, but none have addressed the challenges of accurate inference with multiple linked selected alleles. Many applications search for increases in allele frequencies at sampled sites, but these sites have to be known in advance in order to be sampled [21–23]. Other applications search for local ancestry outliers after applying tools that assume a neutral and uniform admixture process [24–34]. However, selection in an admixed population itself shapes the landscape of local ancestry, and most do not incorporate information about the ancestry tract length distribution. Other approaches have been developed that use summary statistics to detect selection acting in admixed populations, but most do not provide a means to estimate the selection coefficients of the sites under selection [35–39]. Machine learning approaches can be quite powerful for detecting adaptive introgression, but they also usually do not provide a means to estimate selection coefficient, and it is sometimes difficult to interpret the biological underpinnings of the model [40–42]. Our recently developed method resolves some of these difficulties by explicitly modeling selection during admixture in sequence alignment data rather than genotypes as a part of local ancestry inference. Using this approach, it is possible to fit a model with a single locus experiencing additive selection [43], but this has not been generalized for evaluating multi-locus selection. Although a huge range of new methods are rapidly being developed, thus far, none have considered the effects of multiple linked selected sites.

The previous approaches are suitable for finding evidence of selection at a single site in an otherwise neutrally-evolving genome, but in general they do not account for cases where multiple selected sites are genetically linked and may affect the trajectories of each other due to linkage [19,20,43]. Existing methods are expected to incorrectly estimate the selection coefficients of individual sites when they are impacted by linkage with other selected alleles. This makes estimating the selection coefficient of each variant more difficult within admixed populations [43]. These methods also cannot distinguish between single and multiple site selection models, which may lead to an overestimation of the number of selected sites present on a chromosome within an admixed population in some circumstances. We therefore do not have the tools to investigate the impacts of multi-locus selection in admixed populations, but we expect that this phenomenon is widespread for reasons we described above.

We introduce an approach for modeling the effects of linkage between multiple selected sites within admixed populations. We validated our method under a variety of simulated scenarios, where the introgressing population introduced multiple alleles under selection. This approach can accurately identify the number of linked selected sites, as well as determine their location and estimate their selection coefficients by considering the impacts of the linked selected alleles. We applied our method to an admixed population of *D. melanogaster,* and we show that a previous method may have overestimated both the number of selected sites and their selection coefficients due to the effects of linkage between nearby selected alleles. Our results suggest that this is an important contributor to evolutionary outcomes in admixed populations, and our work provides a powerful generalized tool to quantitatively investigate those effects.

## Results and Discussion

### Model Overview

To investigate the impacts of selection on many linked sites, we developed Ancestry-HMM Multi-Locus-Selection (AHMM-MLS). AHMM-MLS is an extension of Ancestry_HMM [44], the latter of which infers both local ancestry and time since admixture for admixed populations by modeling local ancestry, or the ancestry of admixed individuals at particular loci, in a set of samples from the same admixed population as a hidden Markov model (HMM) using a neutral single or multi-pulse admixture model [45]. Our framework considers only single pulse admixture demographic models. The hidden states of the HMM are the local ancestries of the samples at sites across the chromosome, and the observed states are the alleles in the aligned reads at each ancestry informative position. The emission probabilities of the HMM are computed based on the read alignment data from admixed samples and the genotype frequencies in reference unadmixed populations (see [44] for details). AHMM-MLS uses the same emission probabilities as in prior work.

Our method introduces a new technique to infer the expected transition rates between adjacent ancestry informative sites along the chromosome under generalized models of multi-locus selection during admixture in a single pulse model (see below). Because alleles near to the selected sites tend to hitchhike to the selected site, the tract lengths for contiguous regions of local ancestry tend to be longer around selected sites [46], and this effect can be captured in the transition rates between ancestry types. We compute the expected transition rates using a numerical method (see below) and then we use the forward-equations to compute model likelihoods and a direct search algorithm to optimize the model of multi-locus selection.

### Generating Transition Probabilities

The key innovation in this work is the way in which we generate the transition probabilities for the HMM. To capture the effects of multiple selected sites on the transition rates between ancestral states at adjacent ancestry informative sites, we numerically calculate the ancestry states at these sites and at the selected sites after *t* generations. We accomplish this by keeping track of the expected distribution of haplotypes after each generation, numerically calculating the expected effects of recombination and natural selection (see Methods). This allows us to calculate the expected trajectories of haplotypes with many selected sites. Note that our approach does not include the effects of genetic drift and instead assumes the admixed population is infinite. Below we evaluated the impacts of population size on resulting inferences, but as an approximation we expect that drift will have small effects in large admixed populations particularly when admixture occurred recently. Once the transition rates between each pair of adjacent ancestry informative sites are calculated, we use the forward equations to calculate the likelihood of the model given the read pile-up or genotype data from admixed samples. To fit a multi-locus selection model, we use the Nelder-Mead direct search algorithm [47] to optimize the parameters of a multi-locus selection model. These parameters include the location of the selected sites along the chromosome, their selection coefficients, and dominance. We do not consider epistatic interactions among sites in this framework.

### An Iterative Multi-Locus Model Selection Procedure

We developed, tested, and implemented an iterative procedure for fitting a multi-locus selection model to genotype data from admixed populations (Fig 1). Our method begins by identifying a set of potential selected sites using Ancestry_HMM-S [43], and applying basic local-optimum selection approaches to remove trivially close positions that may correspond to or be indistinguishable from a single selected allele (see Methods). We expect that even basic local-ancestry outlier analysis [48,49] might be sufficient for generating a set of candidate positions, but we do not evaluate alternatives here. We then sort the list of candidate selected sites by decreasing likelihood ratio. In the first iteration, we test the site with the highest likelihood ratio as a single site selection model against a null model consisting of a neutral admixture model with similar demographic parameters. In each subsequent iteration, we add one additional selected position, constructing an alternative model that we test against the model obtained from the previous iteration.

**Fig 1.**
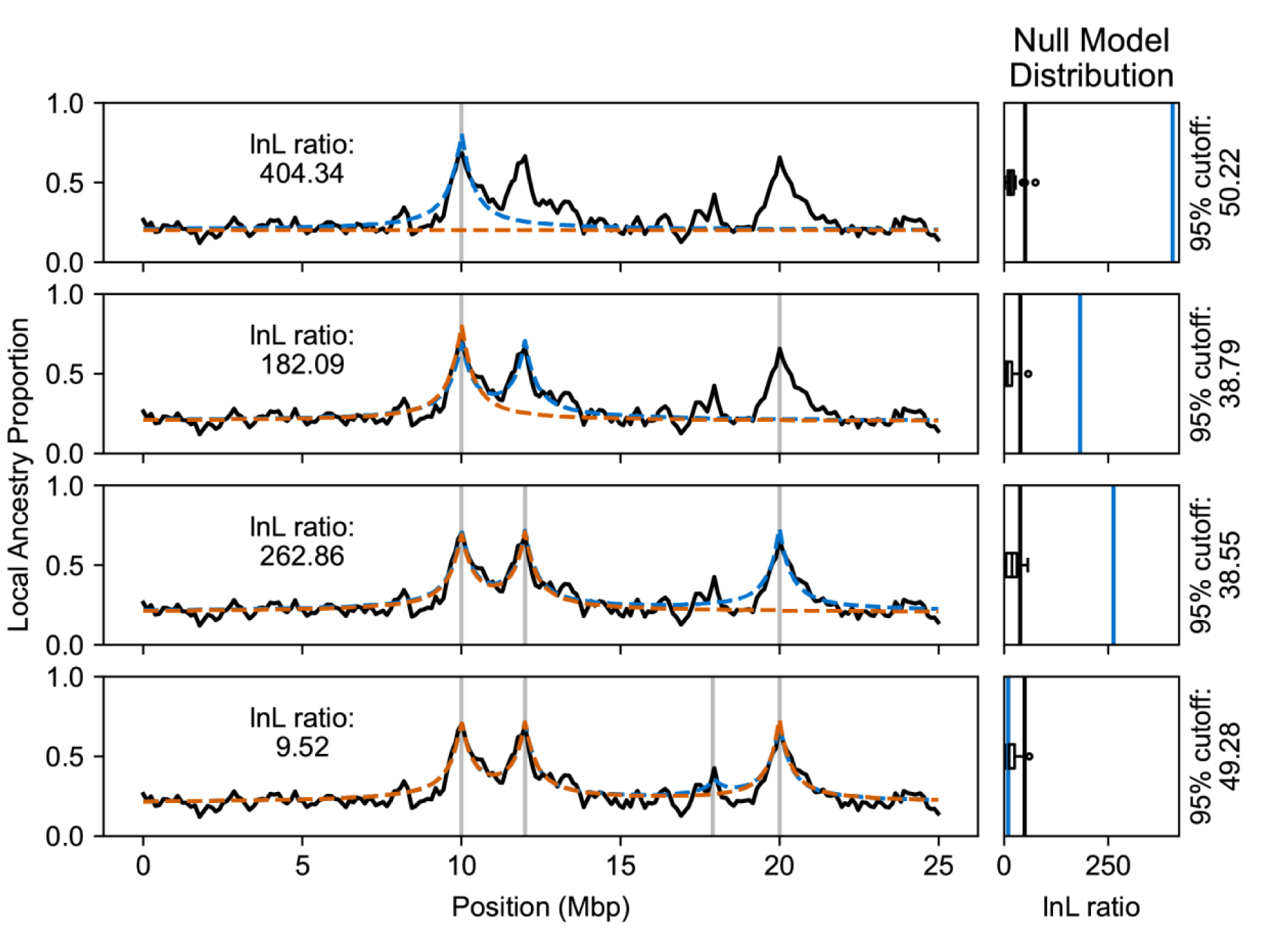
An example of our model selection procedure with four candidate selected positions. On the left, each panel shows the estimated local ancestry proportion within the admixed population (black). On the right, each panel shows the distribution of log likelihood ratios between the alternative and the null model on simulations of the null model (black). In the first iteration (top), we test a single site selection model (blue) against a neutral null model (orange). We then test a two site selection model (blue, second row) against the previous single site selection model which is now the null model (orange, second row). In the third iteration, we test a three site selection model (blue, third row) against the two site selection model from the second iteration (orange, third row). Finally, in testing a four site selection model (blue, bottom), the likelihood ratio (blue, bottom right) does not exceed the 95th percentile of null simulations (black line, bottom right) of the previously selected three site model (orange, bottom). The method terminates and accepts the three site selection model obtained in the third iteration. In each panel, vertical gray lines indicate the positions of selected sites in the alternative model considered.

To obtain the expected distribution of likelihood ratios under the null model, we perform simulations of populations with selections that match this model, and we simulate the sampling of reads from these populations. We do simulations of the null model to obtain this distribution as our method makes assumptions about the admixed population that may not be met. Indeed, simulations under the null model show that population size is a contributor to additional variation not captured by the theoretically expected distribution (S1 Fig), and we expect that other unmodelled components may also affect inferences. On these simulated reads expected from the null model, we fit both the null model, and an alternative model which includes the same sites as the null but additionally includes the site with the next highest peak found by fitting Ancestry_HMM-S. Essentially, evidence for selection at each site is evaluated in the context of all previously estimated selected sites. When an alternative model exceeds the 95th percentile of the simulated null model likelihood ratio distribution, we accept that position and this new multi-locus selection model becomes the null model in the next iteration. If a given position does not exceed the significance threshold, we discard it and attempt the same procedure with the next candidate position against the same null model. The procedure terminates once we have either accepted or rejected every candidate selected site (Fig 1).

### Evaluation of AHMM-MLS Over Simulated Data

In evaluating performance with simulated data (see Methods), we found that our method could accurately distinguish between a single selected site and two nearby selected sites over varying population parameters (Fig 2). We varied the time since admixture *t* from 100 to 1000, the admixture proportion from 0.05 to 0.5, and the distance in Morgans between the selected sites from 0.005 to 0.05. We found that one and two site selection models are easier to distinguish as *t* increases, and as the distance between the sites increases. Our method could also accurately predict the location of the two nearby sites (S2 Fig), when the number of generations is high enough (*t* ≳ 500). The accuracy of the predicted location was not affected by the admixture proportion or as the distance between the sites changes (S2 Fig). Our method could also distinguish between dominant selective pressure and additive selective pressure in certain admixture and selection scenarios (S3 Fig), although this is much more sensitive to the demographic parameters. In most cases, our method performed better as the time since admixture increased, and as the strength of selection increased.

**Fig 2.**
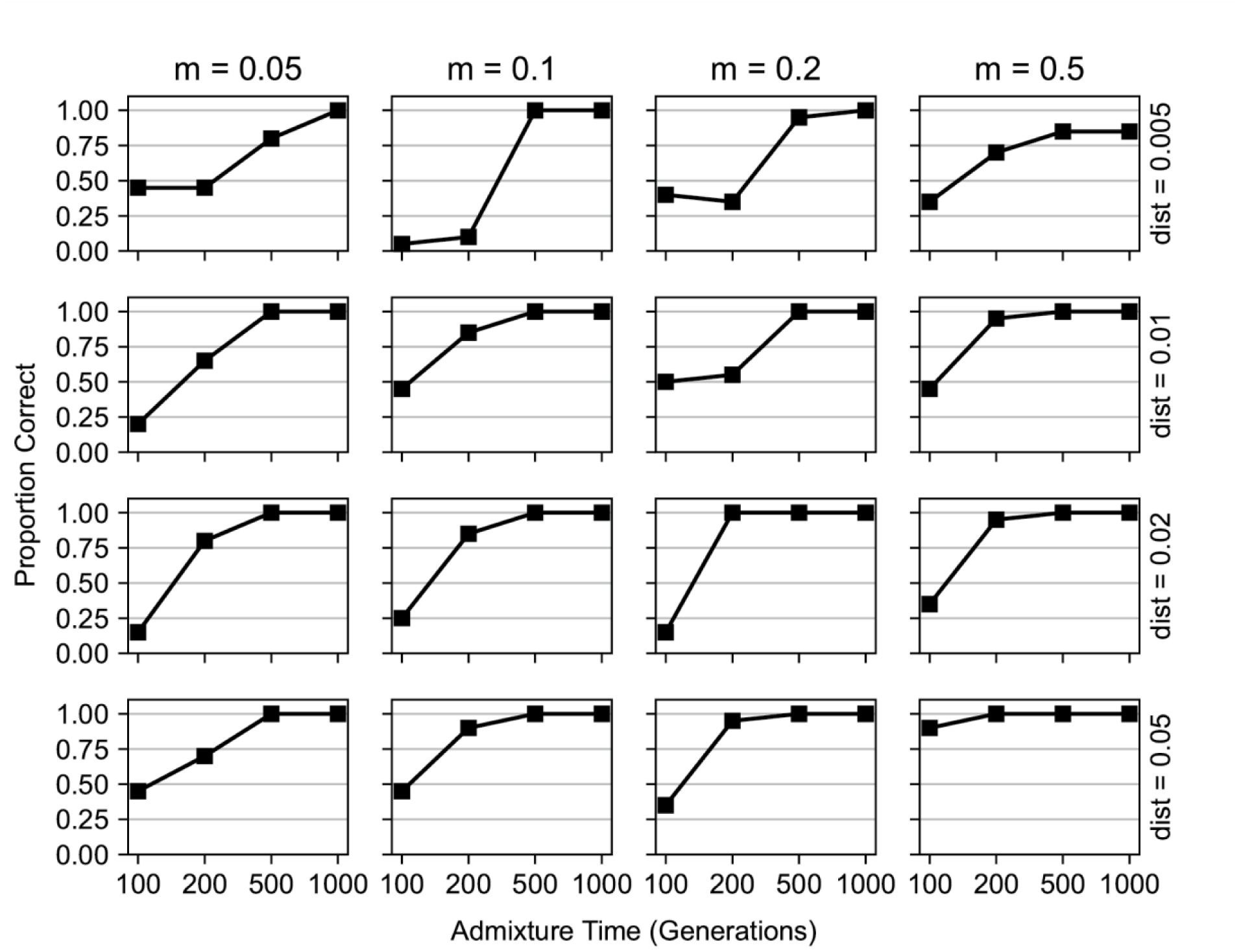
Performance of our method in detecting two nearby selected sites. We evaluated AHMM-MLS in its ability to distinguish between the presence of two nearby sites under selection and the presence of a single site under selection. We ran simulations with varying minor ancestry fractions (0.05, 0.1, 0.2, and 0.5 from left to right), times since admixture (100-1000 generations), and distances between selected positions (0.005, 0.01, 0.02 and 0.05 Morgans, from top to bottom). In each simulation, the selection coefficient of both sites was 0.01. There were a total of 64 different combinations of demographic and selection model parameters. We also ran null model simulations, where there was only a single site under selection introduced in the admixture event, to establish a null model distribution (see Methods). The points on the lines indicate the proportion of two site simulations in which the single site null model was correctly rejected.

We found that optimizing in the correct model space (*e.g.,* with the correct number of selected alleles) gave more accurate predictions of the selection coefficients and of the positions of the sites under selection. For these simulations we varied the selection coefficients of the selected sites *s* from 0.005 to 0.05, and the distance in Morgans between the selected sites from 0.005 and 0.05. In most simulated scenarios, the selection coefficient was considerably overestimated if only a single selected site was fit, as the effects of linkage between selected sites are not considered. This effect was most prominent when the two simulated sites were closer to each other and when their selection was strong (Fig 3).

**Fig 3.**
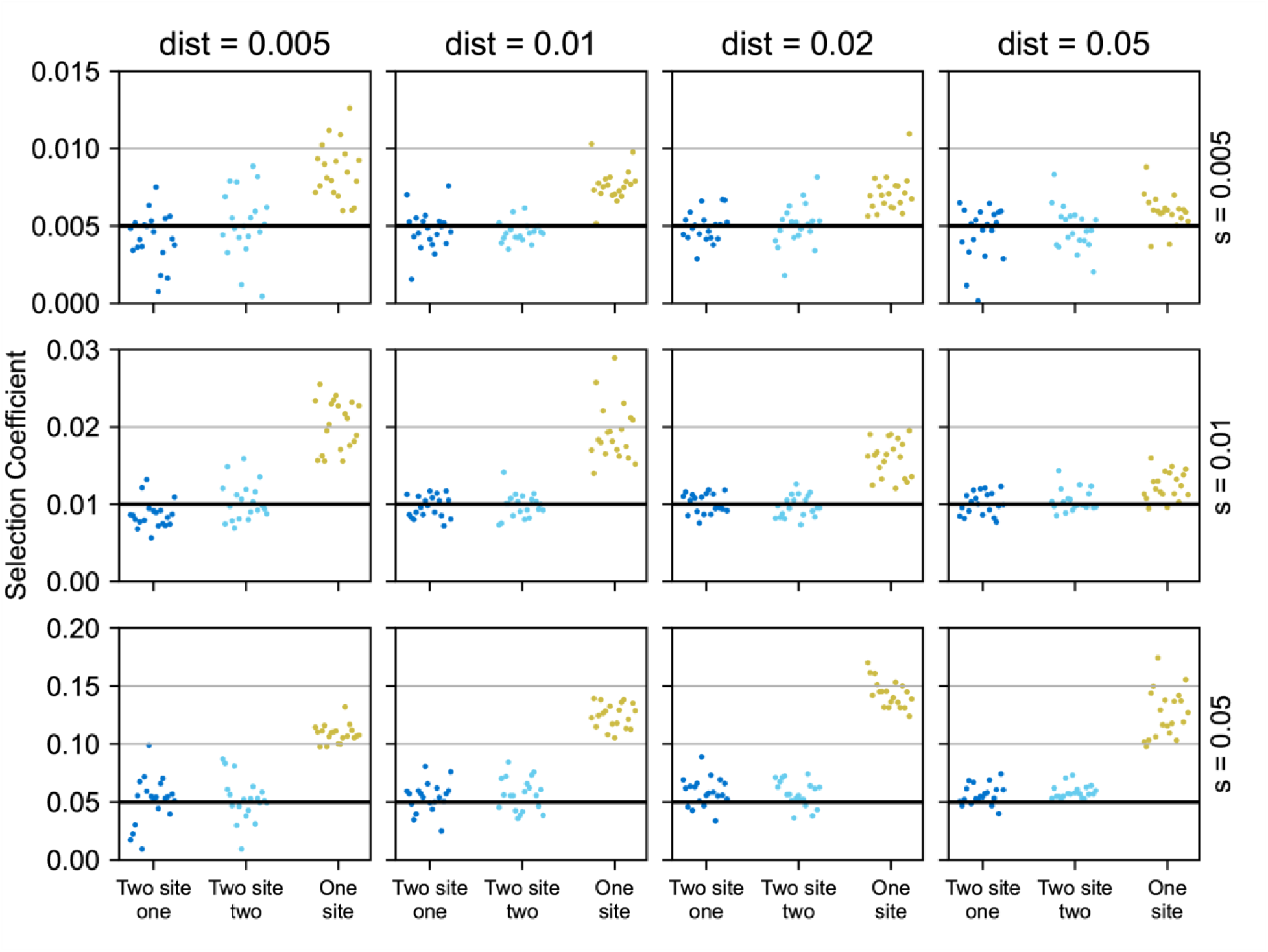
Comparing the inferred selection coefficients between single site and two site models. We evaluated the estimated selection coefficients when using two different models to approach the same data. We ran simulations with varying selection coefficients (0.005, 0.01, and 0.05 from top to bottom) and distances between selected positions (0.005, 0.01, 0.02 and 0.05 Morgans, from left to right). On each simulation we fit a two site model and a single site model. The dark blue and the light blue dots indicate the inferred selection coefficients of the two sites in the two site model, while the yellow dots are the inferred selection coefficients of the single site model. The horizontal black lines indicate the simulated selection coefficients.

We also obtained more accurate estimated selection coefficients in simulations including dominance when we model that site as having dominant, rather than additive, fitness effects (S4 Fig). For relatively recent admixture events (t = 100) and lower selection coefficients (s ≤ 0.02), this effect was most prominent, with the selection coefficients overestimated by nearly 100% when optimizing an additive model on simulations of a site under dominant selection. However, our method generally performs poorly in estimating dominance coefficients (S3 Fig) and we caution against strong interpretation of results obtained from this approach.

Our method could accurately determine the number of linked sites under selection (Fig 4). In simulations, we varied the number of sites to be tested (3 to 4), the distance between the sites (1 to 2 centimorgans), the selection coefficients of the introduced sites (s=0.005 to s=0.01) and the number of generations since the admixture pulse (t=200 to t=500). The simulated populations were the result of an introgression event where the minor population introduced multiple alleles spaced 1 to 2 centimorgans apart undergoing positive additive selection. For simulations where the time since the admixture pulse was 500 generations, we could reliably estimate the correct number of sites. This implies that our method will be appropriate for many admixed populations including populations of sub-Saharan *Drosophila melanogaster* that we consider below, but that caution is warranted for application to recently admixed populations with moderately weak selection.

**Fig 4.**
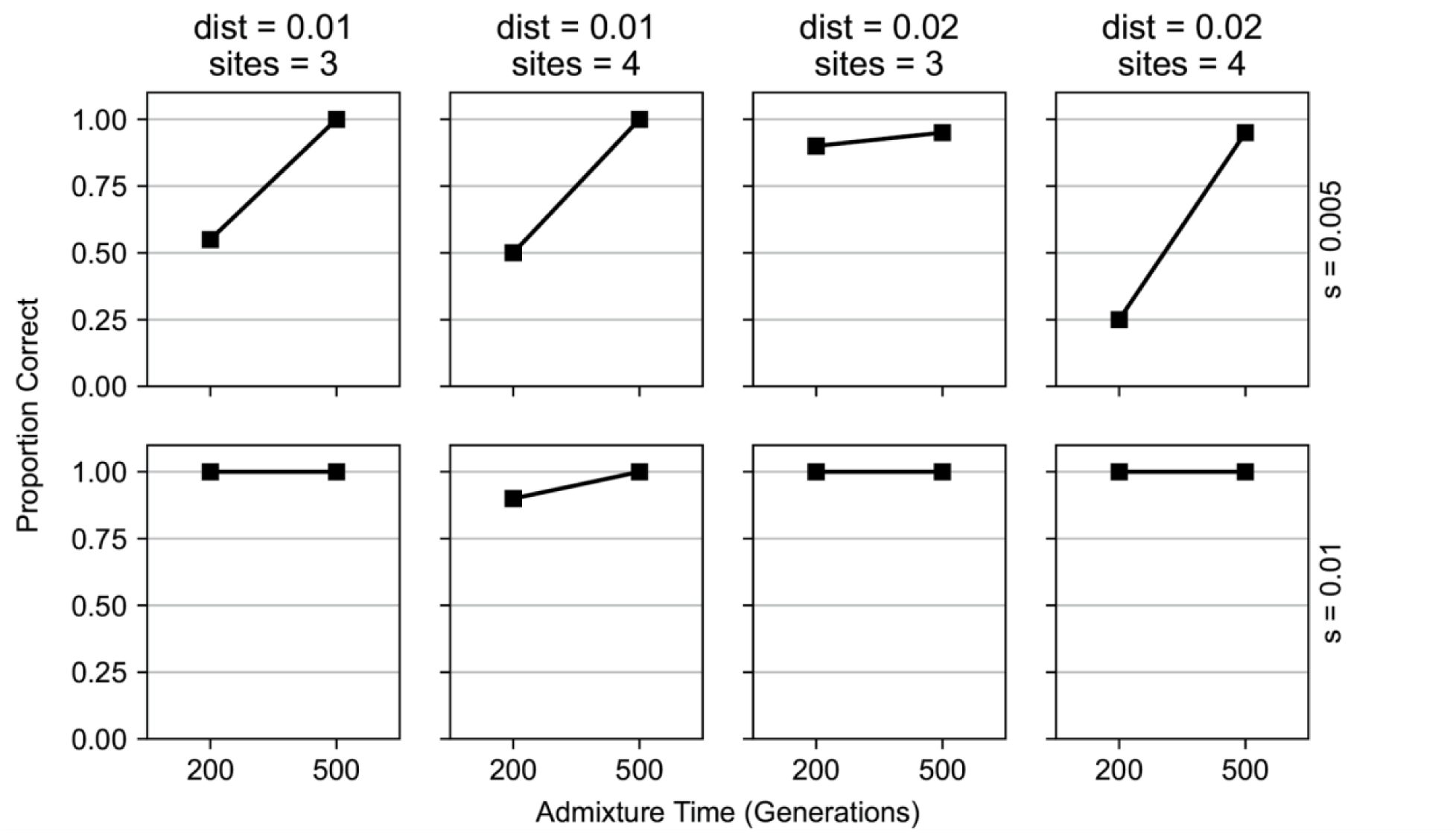
Performance of our method in detecting multiple nearby sites under selection. We evaluated the ability of AHMM-MLS to distinguish 3 site and 4 site models from models with one fewer site. We ran simulations with varying selection coefficients (0.005, and 0.01 from top to bottom), distances between selected positions (0.01 Morgans for the two columns on the left, and 0.02 Morgans for the two panels on the right), and number of introgressed selected sites (3 for the first and third columns, 4 for the second and fourth columns). For the eight different scenarios, we simulated both null and alternative models, where the null model had one fewer selected site. The points on the lines indicate the proportion of alternative model simulations in which the null model was correctly rejected.

### Application to Admixed Drosophila Melanogaster

In order to investigate the potential impacts of interference between linked sites on inferences of natural selection in real data, we applied our method to chromosome 3R of an admixed population of *Drosophila melanogaster* from South Africa. This population shows signals of admixture which have been noted in previous studies [44,45,50]. The admixture history is consistent with a one-pulse model, with admixture parameters that suggest that this population is suitable for our program [45]. In a previous study, we found evidence that chromosome 3R may contain multiple nearby selected alleles [43]. This study had found 13 putative sites under selection on 3R, most of which were fewer than 5 centimorgans away from another selected site. Here we fit models with multiple nearby selected positions, to determine whether and to what extent interference may have impacted our prior estimates of the number of selected sites and their selection coefficients.

### Obtaining A Demographic Model

We first used AHMM [44] on chromosome arm 3L of this population of *D. melanogaster*, to estimate a demographic model for this population. This chromosome arm showed no evidence for the presence of alleles that experienced strong positive selection during admixture [43]. We inferred the admixture fraction and admixture time, obtaining values (*m* = 0.138 and *t* = 466) similar to what we obtained in prior work [45], but we note that here the admixture fraction is slightly lower and the number of generations since admixture is slightly higher. Both differences are consistent with the notion that the presence of selected positions along other chromosome arms may have slightly impacted our prior demographic modeling efforts [44]. This supplied an estimated demographic history that we used as a baseline in our models of selection.

### Identifying Candidate Selected Positions

We ran AHMM-S [43] to evaluate evidence for positive selection along chromosome arm 3R (Fig 5). This program evaluates a neutral model and optimizes an additive selection model and outputs the likelihoods for each. At each position, we recorded the log likelihood ratio between each model. To identify local optima, we performed a simple peak finding algorithm, where we recorded each site that had the highest maximum likelihood of the nearest 1,400 sampled sites and a log likelihood ratio above 15. This gave us 17 candidate selected positions to examine which we ordered by decreasing likelihood ratio. We then applied the iterative procedure described above to produce a multi-locus selection model.

**Fig 5.**
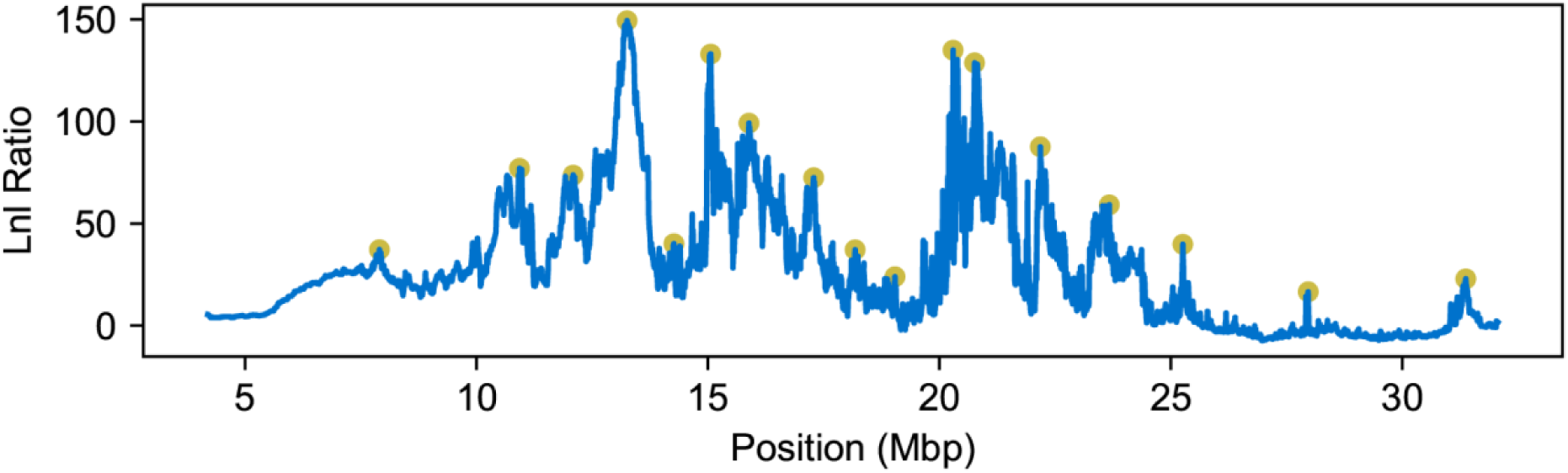
Candidate selected positions on chromosome 3R of *D. melanogaster*. Chromosome 3R of *D. melanogaster* shows signs of many nearby selected sites that are in close linkage. The likelihood ratio outputs of AHMM-S, which test each site for additive selection using a single selected site model, indicate high variation in models of natural selection (blue). Using a simple peak finding algorithm, we identified 17 sites that may be experiencing selection in this admixed population (yellow).

After applying the procedure to iteratively construct a model of multiple selected sites for chromosome arm 3R, we identified 9 selected sites (Fig 6). Comparing this to our previous approach that treated each site as a separate hypothesis test, we found that selection coefficients on 3R may be overestimated. Indeed, the selection coefficients estimated by AHMM-S were up to 49% higher than those found by our method (Table 1). Presumably this occurs because each positively selected site is hitchhiking to an extent on linked positively selected sites, thereby increasing their frequencies in aggregate to a larger degree than would be expected given the selection coefficient at an isolated selected position. This effect would also confound other methods to estimate the selection coefficient, such as those based on excess of local ancestry. We therefore conclude that interference among selected positions has likely impacted analyses of selection during admixture. More generally, the tool we present in this work is a powerful approach for disentangling these potentially complex effects.

**Fig 6.**
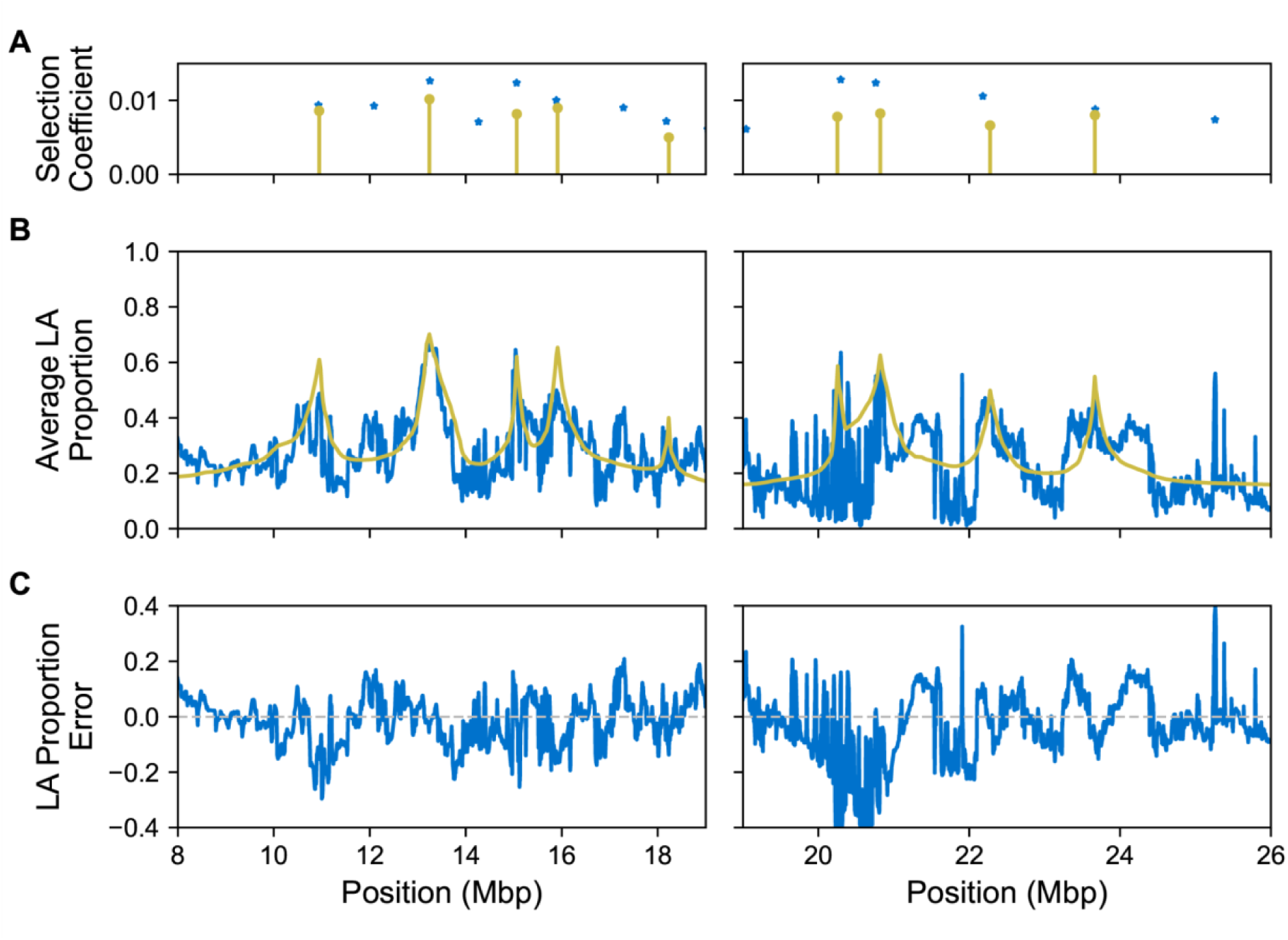
The local ancestry (LA) expected from the model roughly follows the estimated local ancestry of the samples. Due to our method having a time complexity that is exponential with respect to the number of sites, we only calculate these expected local ancestries for half of the chromosome arm at a time (subdivided on the left and right). **(A)** The identified sites and their selection coefficients from our method (yellow) along with 14 out the 17 candidate selected positions from AHMM-S (blue). **(B)** The mean local ancestries from the samples (blue) and the local ancestries expected from the model (yellow). **(C)** Error between the model’s predicted local ancestry and what we infer from the empirical data.

**Table 1.** Comparison of selection coefficients inferred for selected sites by AHMM-MLS and AHMM-S.

Comparison of selection coefficients inferred for selected sites on 3R of South African *D.*
| Approximate<br>base pair<br>position of<br>selected site | Selection<br>Coefficient<br>estimated by<br>AHMM-MLS | Selection<br>Coefficient<br>estimated by<br>AHMM-S | Percentage<br>change | AHMM-S<br>likelihood<br>ratio | AHMM-MLS<br>Likelihood<br>ratio with<br>respect to<br>null model |
| --- | --- | --- | --- | --- | --- |
| 7902645 | Not supported | 0.0073 |  | 37.22 | 8.17 |
| 10946217 | 0.0086 | 0.0093 | 8% | 72.82 | 54.09 |
| 12088137 | Not supported | 0.0093 |  | 73.74 | 6.31 |
| 13239296 | 0.0102 | 0.0125 | 23% | 144.50 | 97.56 |
| 14265104 | Not supported | 0.0071 |  | 40.13 | 0.03 |
| 15062567 | 0.0082 | 0.0122 | 49% | 130.89 | 82.73 |
| 15914622 | 0.0090 | 0.0098 | 9% | 93.24 | 56.21 |
| 17284518 | Not supported | 0.0090 |  | 72.46 | 23.26 |
| 18229524 | 0.0050 | 0.0056 | 12% | 21.92 | 21.46 |
| 19040446 | Not supported | 0.0061 |  | 23.95 | 6.82 |
| 20255076 | 0.0078 | 0.0116 | 49% | 89.97 | 107.26 |
| 20820785 | 0.0082 | 0.0122 | 48% | 116.61 | 39.90 |
| 22277138 | 0.0066 | 0.0094 | 41% | 71.90 | 31.61 |
| 23665629 | 0.0080 | 0.0088 | 10% | 57.31 | 34.14 |
| 25258231 | Not supported | 0.0074 |  | 39.85 | 18.99 |
| 27968043 | Not supported | 0.0053 |  | 16.56 | 0.52 |
| 31371682 | Not supported | 0.0056 |  | 22.90 | 5.53 |

Additionally, we found that many positions that when analyzed individually are consistent with positive selection on admixed ancestry are not supported in the final model. Of the original 17 candidate sites we identified using AHMM-S, 8 were not included in the final model selected by our iterative procedure. As might be expected, these 8 positions are primarily those with lower selection coefficients as estimated by AHMM-S (Table 1). However, when analyzed individually using AHMM-S, we found that each had a large likelihood ratio when comparing a single site selection model to a neutral model (lnL 16.56 - 73.74, Table 1). Support for sites that are relatively distant from other selected positions, such as the candidate site at 31371682 base pairs, were still impacted by the effects of linkage. This result emphasizes the importance of evaluating multi-locus selection models to capture the evolutionary dynamics of natural selection in admixed populations.

### Caveats

Although our method provides a promising means for quantifying and investigating the impacts of interference and natural selection in admixed populations, there are several important caveats. First, the state space of multi-locus selection models is extraordinarily complex, and there is no evaluation procedure that could exhaustively attempt all possibilities. For example, even in the scenario that we considered with 17 candidate selected positions, there are *2^17^*(∼130,000) possible combinations of sites in multi-locus selection models. This is an intractable number of models to test, even without considering the difficulty of estimating selection coefficients. The iterative procedure we present is an appealing way to prioritize model space and we expect that it will perform well in a variety of scenarios, but undoubtedly there are other plausible models that we could not evaluate. Second, our approach will accommodate scenarios where there are a modest set of loci of relatively large effect. However, some authors have proposed that the aggregate effect of hundreds of weakly selected linked mutations might shape the landscape of admixed ancestry in natural populations [9]. Our approach is not well suited to such scenarios because each site is unlikely to reach significance in itself and because the time taken to compute the expected transitions for each model is exponential with respect to the number of sites. We expect that AHMM-MLS will typically perform best with populations that are only somewhat genetically divergent and where strong selection strongly affects the dynamics of introduced alleles.

## Conclusion

Admixture has the potential to simultaneously introduce multiple linked selected sites, but this phenomenon is rarely addressed in empirical investigations. To meet this need, we created AHMM-MLS. In validating our method over simulated data, we found that it could identify multiple nearby selected sites, and estimate the selection coefficients better when the linkage between these sites was accounted for. We found that previous studies of adaptive introgression on chromosome arm 3R of an admixed *D*. *melanogaster* population overestimated the number and strength of selected sites along the chromosome. Because divergent populations may introduce many selected alleles at once, analyzing the effects of linkage between these sites is critical for understanding the evolutionary dynamics of admixed populations. We hope that our method can be applied to the many examples of adaptive introgression that have already been identified, and can better quantify cases where multiple advantageous sites have been introduced at once.

## Materials and Methods

### Calculating Expected Haplotype Frequencies

To calculate the effects of multiple selected sites on the expected ancestry transition rates between two adjacent ancestry informative sites, we use a numerical method to track the expected distribution of haplotypes after *t* generations, including the local ancestries of the selected sites and the two ancestry informative sites. If there are *n* selected sites, then we track *n* + 2 total positions (i.e., selected positions and ancestry informative sites), leading to *2*^*n*+*2*^ haplotypes. We do not assume the selected sites are sampled as ancestry informative markers in the admixed population and their positions may lie anywhere along the chromosome. The haplotype distribution in the admixed population is modeled by a row vector *H*, which undergoes a transformation in each generation, producing a sequence of vectors *H^0^*… *H*^*t*^, one for each generation. The vector *H*^*g*^ is transformed into the vector *H*^*g*+^*^1^*, representing the expected change in the haplotype distribution from generation *g* to *g +* 1. As a convention for this work, *H_0_*is the frequency of the haplotype consisting of positions all originating from ancestral population 0, (i.e. the ancestral origin of the sites along the chromosome is 00…00), and *H_1_*is the frequency of the haplotype 00…01, and *H_2_*of haplotype 00…10, and so on, counting in a binary fashion. At generation 0, only two haplotypes, *H^0^_0_*and *H^0^_2_*_*n*+_*_2−1_*, have non zero frequencies. That is because we assume a single admixture pulse begins from unadmixed population founders *t* generations before the time of sampling and these represent the ancestral haplotypes. These are the two haplotypes where all sites are of the same ancestry, and their initial values are dictated by the admixture fraction *m*.

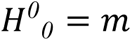

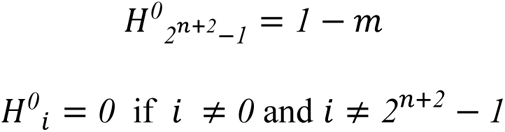

In each generation, we produce a row vector, *D*, which corresponds to the expected distribution of diploid individuals that result from the haplotypes assuming an infinite population, random mating, and no segregation distortion.

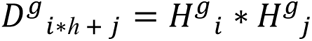

Now, to go from this diploid genotype distribution to the haplotype distribution of the next generation, we apply the matrix operation ***M*** to *D*^*g*^, the result of which is *H′*^*g*+*1*^. *H′*^*g*+*1*^ is then normalized to produce *H*^*g*+^*^1^*.

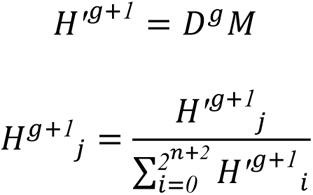

We call the matrix ***M*** the diploid to haploid transformation, as it converts the diploid genotype distributions of one generation to the expected haplotype frequencies of the next generation, accounting for the effects of recombination and natural selection within diploid individuals. A specific entry, such as *M*_*i*,*j*_, is the contribution of the diploid genotype *i* on the haplotype *j*, taking into account the fitness of the diploid genotype *i*, and the probability that recombination events produce the haplotype *j*. ***M*** is constant for each pair of adjacent ancestry informative sites, and so it only needs to be calculated at *t* = 0. The matrix depends on the location of the two adjacent ancestry informative sites, the location of every selected site, and the fitness coefficients of those selected sites. The location is represented as a single value, *l*, the position on the chromosome in Morgans. The two fitness coefficients are the dominance coefficient, *h*, and the selection coefficient, *s*. This would make the relative selection coefficients 1, 1 - *hs*, 1 - *s*. To simplify calculations, we assume all ancestry informative sites — *i.e*. the positions sampled from along the genome in genotype or pileup data — are neutral sites where *s* = 0.

To generate ***M***, we iterate through all possible diploid genotypes. We assume that fitness values combine across selected sites multiplicatively. For a particular diploid genotype *i*, it has an associated fitness *S*_*i*_, which we compute by taking the product of the relevant selection coefficients for each site. For this diploid *i*, we iterate through each region where a recombination event may occur. If there are *n* sites, then there are *n +* 1 regions. Each of these regions has a corresponding recombination rate *r*. Recombination in a specific region would produce two haplotypes, *k* and *l*. So for each recombination event of this diplotype, there would be a contribution proportional to *D*_*i*_ ∗ *S*_*i*_ ∗ *r* to haplotypes *k* and *l*. This contribution is reflected in ***M*** by adding *S*_*i*_ ∗ *r* to the *M*_*i*,*k*_ and *M*_*i*,*l*_ entries. ***M*** is computed after we have iterated through all possible diploid genotypes and added their contributions to each haploid that they may produce through meiosis. By reducing the most computationally expensive parts of the numerical procedure to matrix multiplications, we are able to use the optimized linear algebra library armadillo [51,52] to quickly compute the transition probabilities for each adjacent pair of ancestry informative sites.

After iterating for *t* generations, we are left with the haplotype distribution that is expected under our model of selection assuming an infinite admixed population size. We directly calculate the transition probabilities between the local ancestries of the ancestry informative sites by iterating through the haplotype distribution and recording the rates of the four possible ancestry state combinations for the two ancestry informative sites.

### Model Optimization

Once we have the expected transition rates for a particular model, we use the forward equations to compute the likelihood of this model given the read pile-up or genotype data. We optimize the parameters of the model to maximize its likelihood using Nelder-Mead search simplex optimization algorithm [47]. For our hyperparameter values we used a reflection constant of 1, a contraction constant of 0.5, an expansion constant of 2, and a shrinkage constant of 0.5. For a single site with unrestricted selection, the location of the site and two selection coefficients, *h* and *s*, are optimized. If *h* is known and fixed (*e.g.,* if additive *h* = 0.5), then only one selection parameter needs to be optimized.

For each optimization, an initial starting point for the location of the selected sites must be supplied by the user. Each optimization is done in two stages, where in the first stage the simplex is centered around the supplied starting point, while in the second stage the simplex is centered around the optimum of the first stage. Each stage consists of multiple starts, with different simplex sizes and orientations. If the range of log likelihoods for every point in the simplex falls below a certain threshold, or if four shrinkage transformations occur in a row, then that search is stopped and the optimum of the simplex is taken to be the optimum of that search. In the first stage, the search is stopped if the range falls below 5 and it falls below a quarter of the initial simplex range. In the second stage, the search is stopped if the range falls below 1 and it falls below one 20th of the initial simplex range.

### Likelihood approximation

To reduce the time taken to optimize each proposed model, we only calculate the effects of a selected site on the transition rates between two ancestry informative sites if those sites are less than a specified distance from the selected site (with a default of 2 centimorgans). When this distance is smaller than the distance between two sites, and those sites have a strong selection (*e.g.,* selection coefficient of 0.05), then the selection coefficients for these sites can be overestimated (S5 Fig). We recommend increasing this distance when estimating models with strong selection. We also only calculate the transition rates between every k-th pair of adjacent ancestry informative sites, as an adjustable parameter (default of 4). We found that ignoring all but every k-th pair of adjacent ancestry informative sites does not affect the inference of selection coefficients when k is relatively small. If the local ancestry must be decoded for a given MLS model, then we calculate the effects of sites throughout the entire chromosome. This can be costly, as the size of the matrix ***M*** is exponential with respect to the number of sites on each tracked haplotype, so we don’t recommend doing this for models with more than 6 sites. We instead recommend that users split the model into pieces with sites that are decently separated, as we have done with 3R above.

### Forward simulations

To evaluate our method over a large variety of plausible introgression scenarios, we performed forward simulations. In each of our simulations, a diploid population received a single admixture pulse from another population carrying at least one selected allele, producing a large admixed population (*n* = 10000, unless stated otherwise). As in prior work [43–45], we first used the coalescent simulation program MACS [53] to create the genotype data for unadmixed individuals. We simulated the local ancestry along the genome of admixed samples using SELAM [54]. This procedure is described in detail in prior work [44,45]. For each population and selection scenario, we simulated 20 admixture events that aligned with the null model and 20 that aligned with the alternative model. We sampled 75 diploid individuals from each of these simulations. We evaluated AHMM_MLS on its ability to distinguish between alternative and null models (see below).

### Creating a null distribution

Our method makes simplifying assumptions about the admixed and ancestral populations that may make it unsuitable for common model selection methods such as the Bayesian information criterion and the Akaike information criterion (S1 Fig) [55,56]. Instead, we generate the expected likelihood ratio distribution for a null model by performing several simulations of a population evolving under the null model. We then fit both a null model and an alternative model which usually includes one additional selected site to the simulated data, and record the distribution of likelihood ratios between the two models in every simulated population. We then apply both models to the population being tested, and if the likelihood ratio between these models exceeds 95% of those on the null simulations, then we accept the alternative model.

### Robustness to demographic model misspecification

We ran simulations of two nearby selected sites and, in different simulations, we misspecified both the time since admixture and the admixture fraction by factors of 0.5, 0.8, 1.2, and 2. As the true time since admixture increased, our method was generally more robust to time misspecification. When the true time since admixture was 500 generations or higher, misspecifying by a factor of 0.5 or 2 produced only small effects for the ability of AHMM_MLS to correctly support the alternative model (S6 Fig). The estimated strength of selection was affected when the true time since admixture was small (*t* = 100), with most estimated selection coefficients being off by more than 50% (S7 Fig). This effect decreased as the true time since admixture increases to 500 or 1000, where most estimated selection coefficients had an error less than 30%. When the true admixture fraction was relatively small (m ≤ 0.2), our method was not strongly impacted by misspecifications of the admixture fraction by a factor of 0.5 or 2 (S8 Fig).

### Effect of population size

For relatively moderate or long times since admixture (such as our default 500 generations) and with moderate selection coefficients (s=0.005-0.05), our method starts to perform poorly when the effective population size of the admixed population drops below 5000 (S9 Fig). We ran forward simulations of admixed populations with various numbers of individuals, and found that the ability to distinguish between two site and single site models was severely limited once the number of individuals dropped below 5000 (S9A Fig). The accuracy of the inferred locations and selection coefficients of the two simulated sites were also hampered when the number of individuals was below 5000 (Figs S9B and S9C). This is because our method assumes an infinite population size, with no genetic drift, and when populations are too small and admixture is relatively ancient, this assumption will tend to produce suboptimal results.

### Drosophila data and analysis

We used publicly available datasets of *D*. *melanogaster* collected from South Africa [57]. In a previous study, the data was prepared so that it could be analyzed by the AHMM programs [43]. This included removing the known large chromosomal inversions found on some of the chromosome arms [50,58,59]. We used a publicly available fine-scale recombination map of chromosome 3R [60]. However, we note that differences in the assumed map and the true combination rates in this population may impact the accuracy of inferences we obtained here.

When we applied the iterative procedure (see Results) to chromosome arm 3R of this population, we inferred the selection coefficients of each selected site in each model, while keeping the location of the selected site fixed. Once we identified the 9 selected sites, we reoptimized the selection coefficients and locations of the sites to arrive at our final model.

When performing simulations under the null model to determine the expected likelihood ratio distribution, we used a population size of 10,000. This is probably smaller than the effective population size in natural *D. melanogaster* populations. Presumably, because smaller populations are more impacted by drift, we expect that this will be a conservative choice when evaluating evidence for positive selection in this natural population (S1 Fig). If the true population size is larger in reality, additional candidate positions might exceed the significance threshold, but the selection coefficients are not expected to change substantially.

### External tools and libraries

We used the Armadillo c++ linear algebra library to perform matrix multiplication [51,52]. We used GNU parallel to run many batches of simulations at once [61]. We used the FlyBase sequence coordinates converter to convert assembly 5 base pair coordinates to assembly 6 [62].

### Data and code availability

The source code for Ancestry_HMM-MLS is made available at <u>github.com/genicos/ahmm_mls</u>. The scripts used to perform the iterative method on chromosome 3R are available at <u>github.com/genicos/ahmmmls_iterative_site_testing</u>. The scripts used to generate the figures present in this paper as well as the underlying data are available at <u>github.com/genicos/ahmmmls_graphs</u>.

## Acknowledgments

We thank all members of the Corbett-Detig lab. We thank in particular Jesper Svedberg, Jakob McBroome, and Rasmus Nielsen for helpful feedback and discussion.

---

**S1 Fig.**
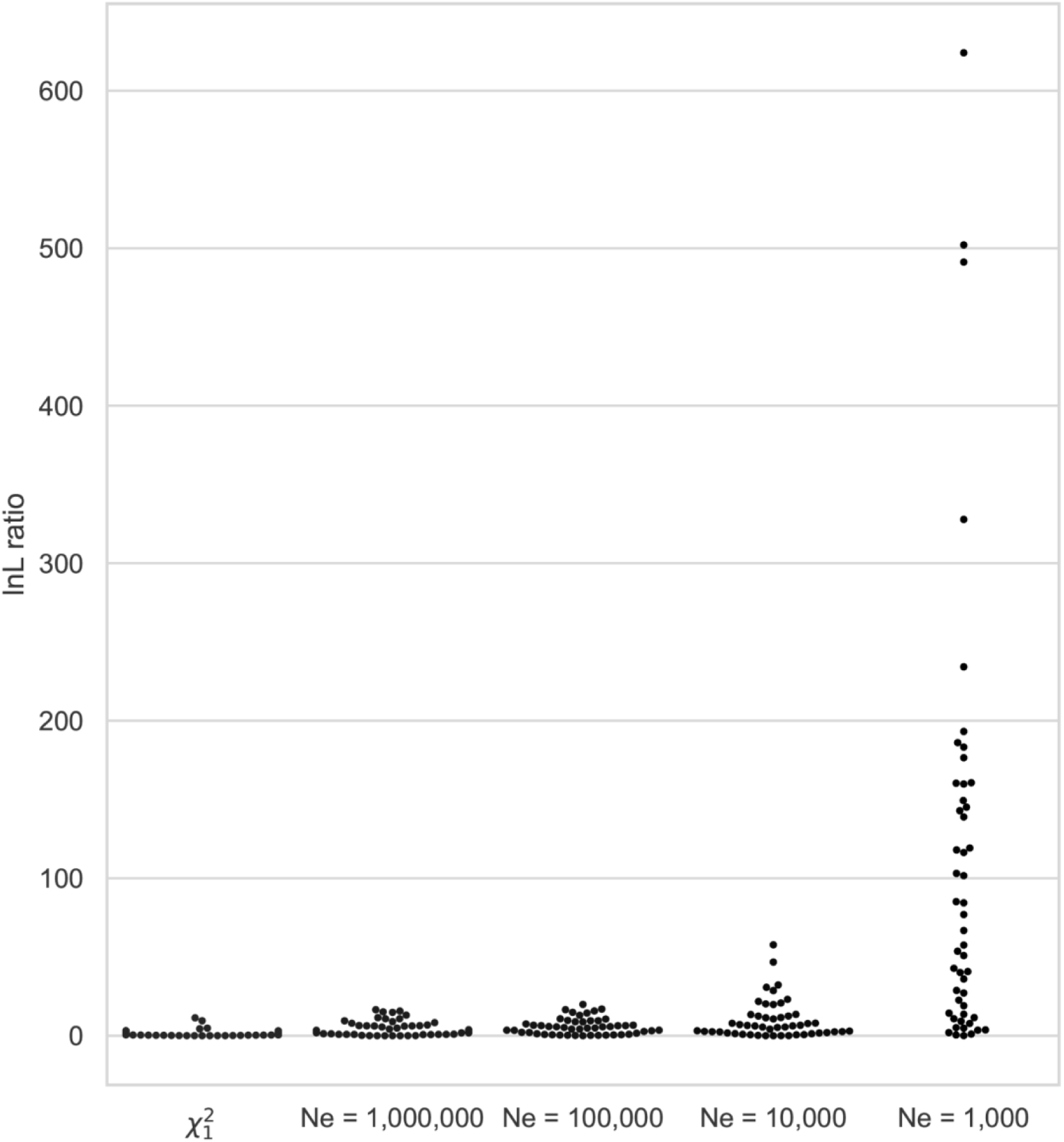
Finite populations skew the likelihood ratio distributions. We simulated 50 neutral admixed populations with admixture parameters *m=*0.2, *t*=500, and varying population sizes. On each simulation we fit a neutral model and we fit the selection coefficient of a single selected site with the location and dominance coefficient fixed. The swarm plots show the log likelihood ratio between these two models, as well as the theoretically expected chi-squared distribution when fitting a single parameter. Our model assumes an infinite population, so as the simulated population grows larger, the log likelihood ratio distribution more closely matches the theoretically expected distribution. We note that the finite population size does not fully account for the disconnect between the simulated distributions and the theoretical distribution.

**S2 Fig.**
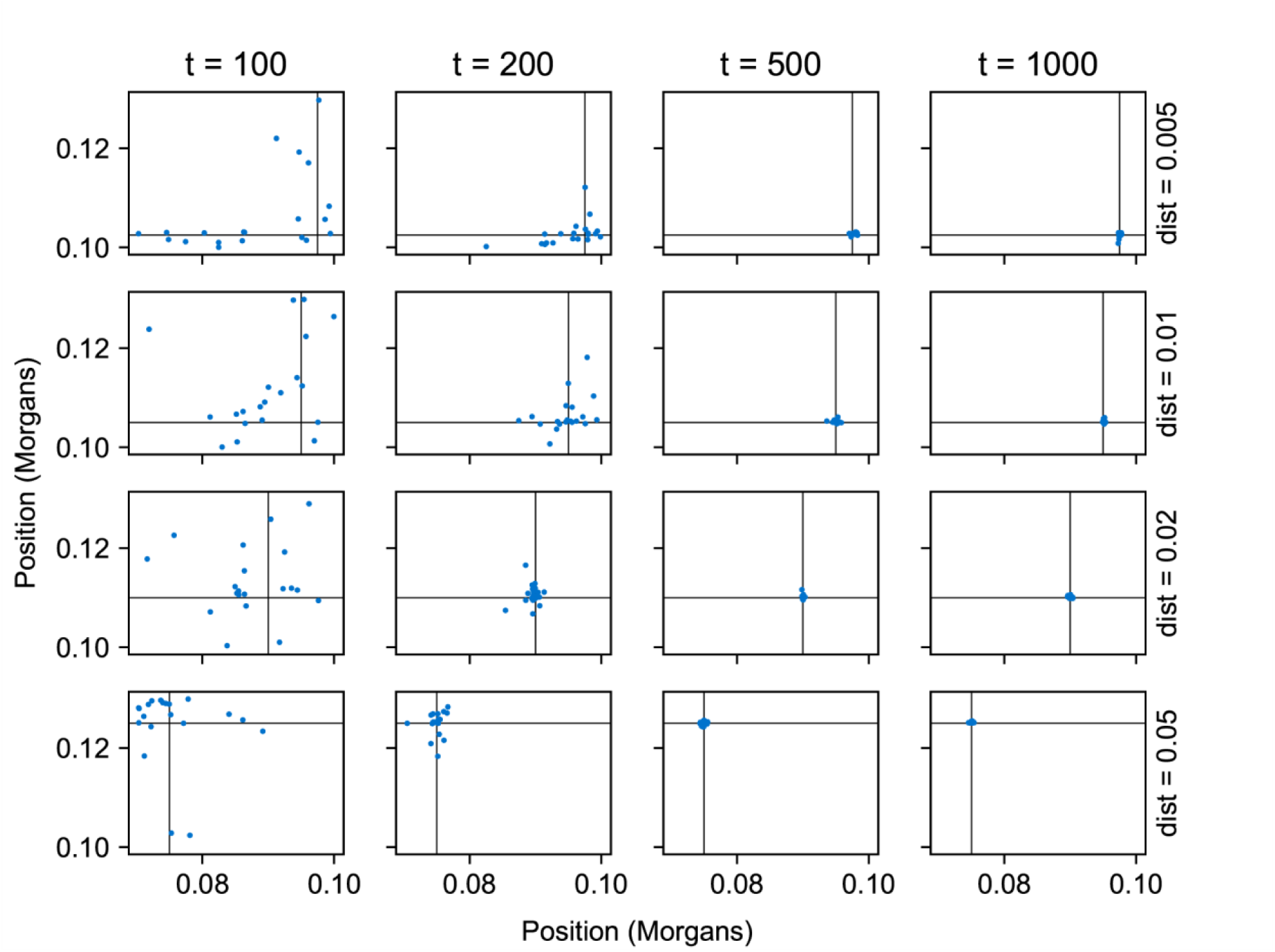
Accuracy of the estimated positions of two selected sites. Each simulated chromosome was the result of an admixture event where *m* = 0.2, and the minor population introduced two nearby sites under positive additive selection with a selection coefficient of 0.01. For each of the 16 graphs, we ran 20 simulations which share the same distance between selected sites (0.005, 0.01, 0.02 and 0.05 Morgans, from top to bottom) and time since the admixture pulse (100, 200, 500, and 1000 generations). On the x-axis and y-axis are the locations of the selected sites on the chromosome in Morgans. The black lines going through the graphs are the true locations of the simulated sites.

**S3 Fig.**
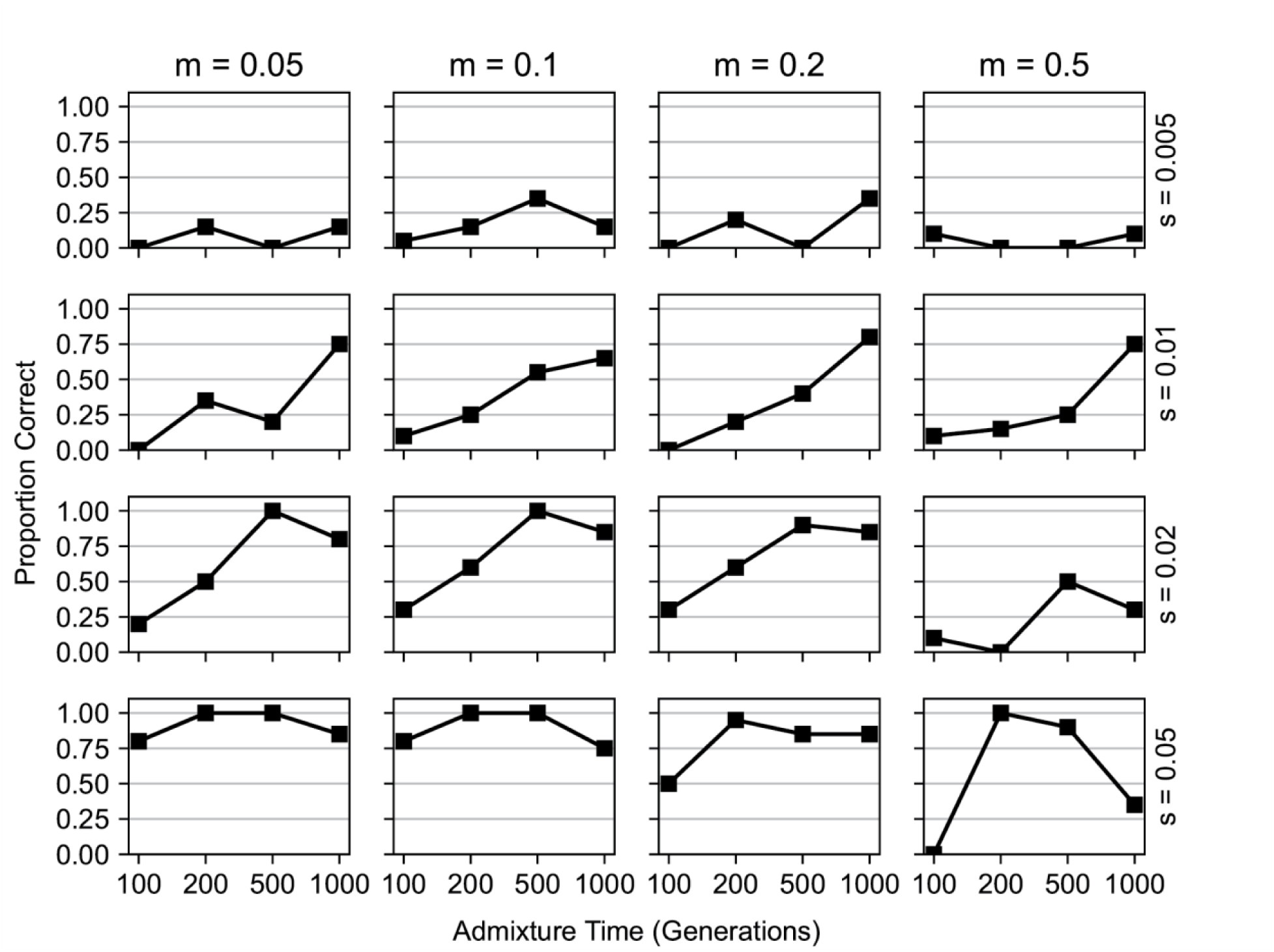
For strongly selected positions, AHMM_MLS can distinguish between selected sites under dominant vs additive selection. Much like the simulations with two sites, we simulated 64 different introgression and selection scenarios, in which the introgressing population contributed a positively selected allele. We varied the minor ancestry fractions (0.05, 0.1, 0.2, and 0.5 from left to right), times since admixture (100-1000 generations), and selection coefficients (0.005, 0.01, 0.02 and 0.05, from top to bottom). For each introgression scenario, we ran null model simulations where the site had additive selection, and alternative model simulations where the site had dominant selection. The points on the line show the proportion of alternative model simulations in which the null model was correctly rejected (see Methods).

**S4 Fig.**
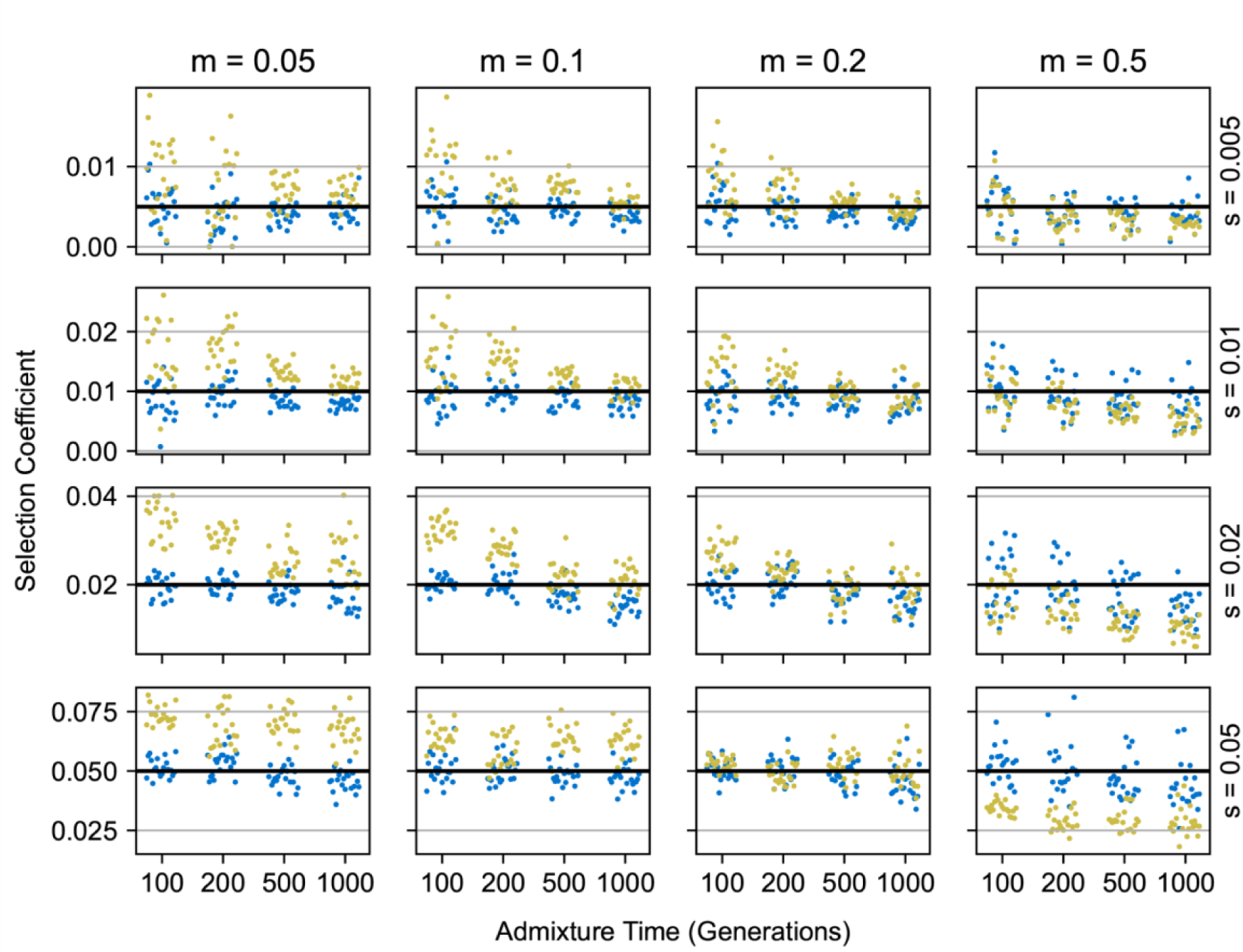
Fitting a dominant selection model on simulations of dominant selection gives more accurate inferred selection coefficients. We inferred the selection coefficients of selected sites for the same simulations from supplemental figure 3. The selection coefficients inferred when fitting a dominant model (blue, h = 1) are much closer to the simulated value (black line) than those coefficients inferred when fitting an additive model (yellow).

**S5 Fig.**
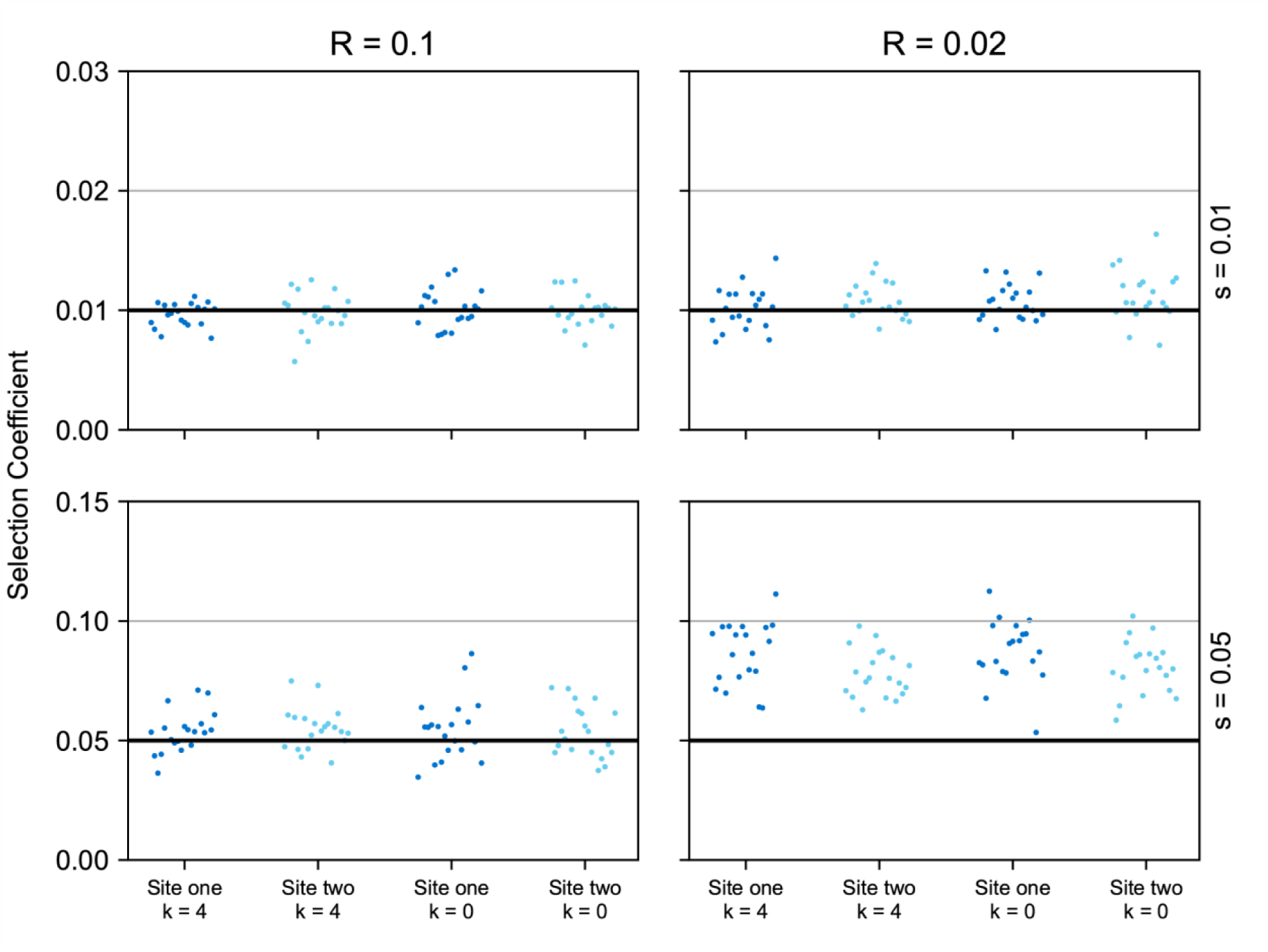
Speed ups employed in our method can affect the inference of strong selected sites that are far away. We simulated an admixed population with two sites under selection, 0.05 Morgans apart, in which both had a selection coefficient of either 0.01 (top panels), or 0.05 (bottom panels). On these simulations we fit a two site model with different speed up parameters in place. In light and dark blue are the inferred selection coefficients, and the black lines indicate the simulated selection coefficients. We altered the number of pairs of adjacent sampled sites that we skipped in the transition rate calculation, *k*, and found that it had very little effect. We also altered the radius around each site in the model where we account for the effect of that site (R=0.1 Morgans, on the left panels, and R=0.02, on the right panels) and found that for large selection coefficients a radius large enough to encompass both sites is required for accurate inference.

**S6 Fig.**
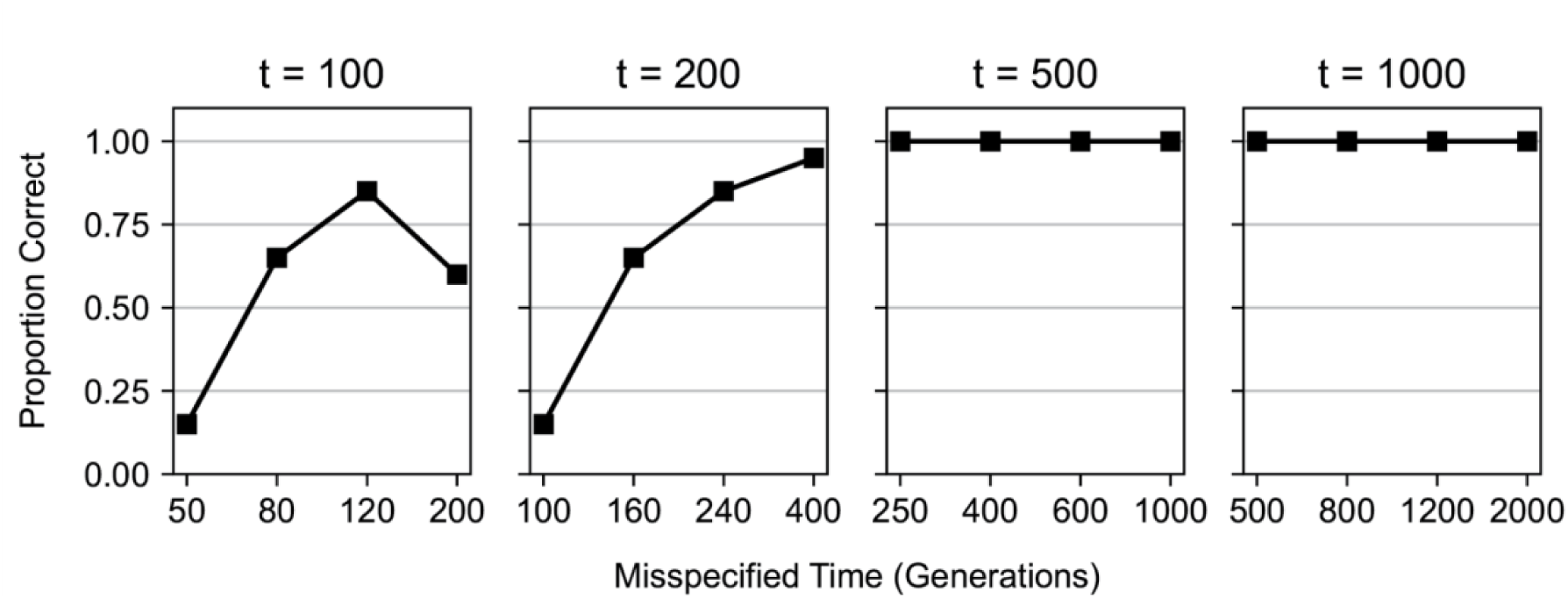
Evaluating the effects of misspecifying the time since admixture when comparing two site and single site models. We misspecified the time since admixture by a certain factor from the true simulated time when analyzing simulations with a single site under additive selection or two sites under additive selection. For the simulations with two selected sites, they were placed one centimorgan apart. In every simulation, the sites had a selection coefficient of 0.01, and an admixture proportion of *m* = 0.2 . We varied the time since admixture from 100 to 1000 generations since admixture (left to right), and misspecified this time in our AHMM_MLS models by a factor of 0.5 to 2. The points on the lines indicate the proportion of two site simulations in which the single site null model was correctly rejected.

**S7 Fig.**
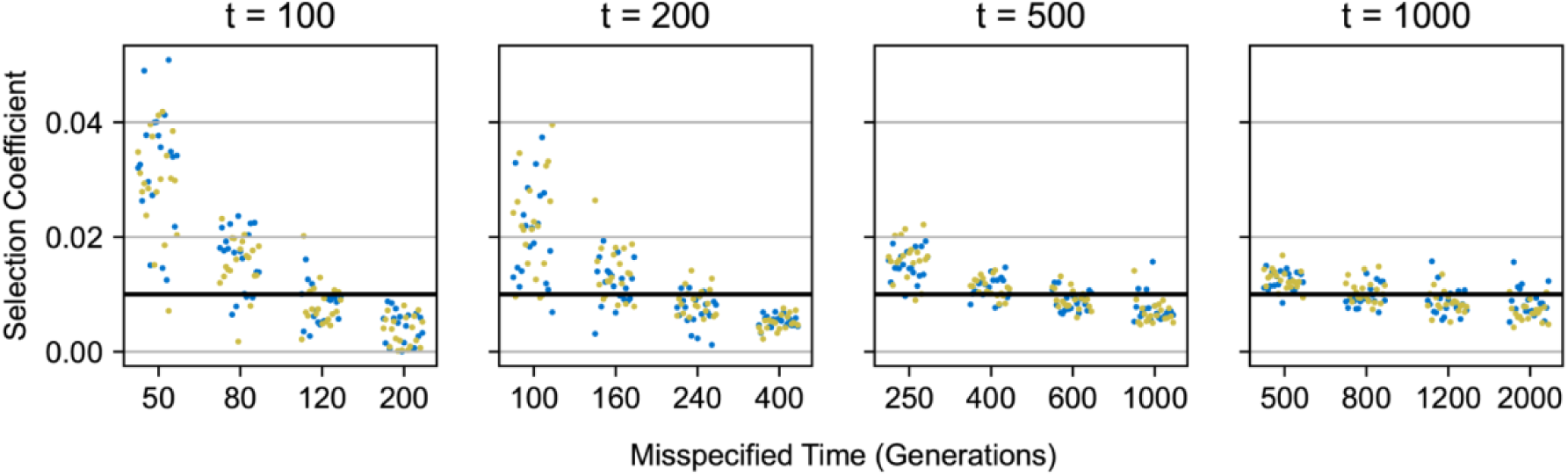
Comparing the inferred selection coefficients when misspecifying the time since admixture. We compared the inferred selection coefficient versus the simulated selection coefficients for the two site simulations from supplemental figure 6. In blue we show the inferred selection coefficients for one of the two sites, and in yellow we show the other. The black line indicates the simulated selection coefficients of both sites.

**S8 Fig.**
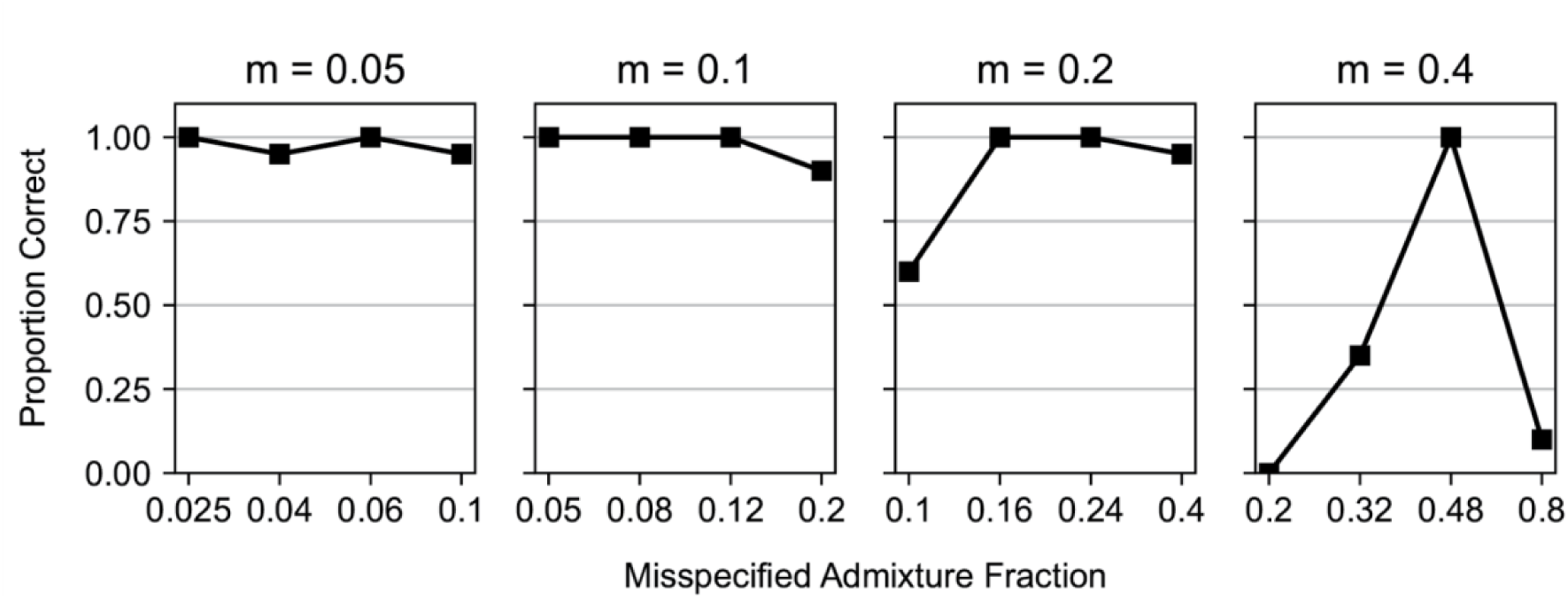
Evaluating the effects of misspecifying the admixture fraction when comparing two site and single site models. We misspecified the admixture fraction by a certain factor from the true simulated fraction when analyzing simulations with a single site under additive selection or two sites under additive selection. For the simulations with two selected sites, they were placed one centimorgan apart. In every simulation, the sites had a selection coefficient of 0.01, and the time since admixture was 400 generations. We varied the admixture fraction from 0.05 to 0.4, and misspecified this fraction in our AHMM_MLS models by a factor of 0.5 to 2. The points on the lines indicate the proportion of two site simulations in which the single site null model was correctly rejected.

**S9 Fig.**
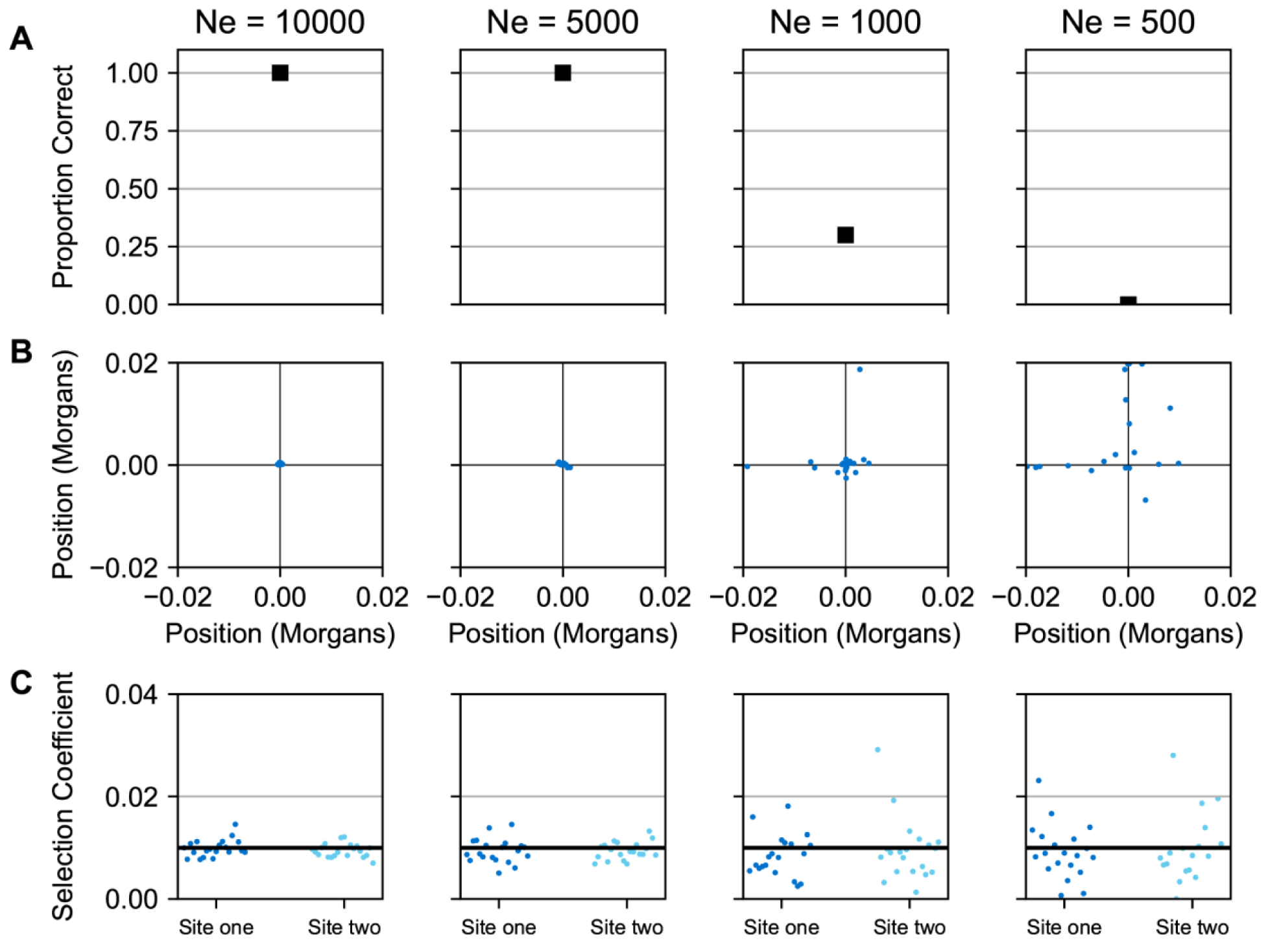
Evaluating the effects of population size. We simulated populations with admixture parameters *m* = 0.2 and *t* = 500, in which the introgressing population brought in two selected alleles with selection coefficients of 0.01. We also simulated null model cases in which only a single selected allele with the same selection coefficient was introgressed. For each of these population models, we simulated populations with varying numbers of individuals, (Ne = 10000, 5000, 1000, and 500 from left to right). **(A)** The proportion of two site simulations in which the number of sites was correctly estimated. **(B)** The position in Morgans of the two inferred selected sites (blue) and the simulated positions (black lines). The x-axis is the position of the first selected position, and the y-axis is the position of the second selected position. Each blue dot corresponds to fitting two sites on a single simulation. Both axes have been translated so that the simulated position is at 0 Morgans. **(C)** The inferred selection coefficients of the two sites (dark and light blue), and the simulated selection coefficients (black).

## Notes

### Competing Interest Statement

The authors have declared no competing interest.

